# Reactive astrocyte derived extracellular vesicles promote functional repair post stroke

**DOI:** 10.1101/2022.09.06.506818

**Authors:** Shangjing Xin, Lucy Zhang, Nhi V. Phan, S. Thomas Carmichael, Tatiana Segura

## Abstract

Reactive astrocytes are both neurotoxic and pro-regenerative. Their reparative roles after injury have been demonstrated, but how they play a contributing role to regeneration remains question. Here, we investigate the use of astrocytic extracellular vesicles from primary astrocytes cultured in reactive conditions in promoting repair after ischemic stroke. Our studies show that extracellular vesicles derived from reactive astrocytes that co-express a significant number of reactive genes (155 upregulated including log2 of 9.61 for *Lcn2*) and axonal outgrowth genes (59 upregulated including log2 of 3.49 *Ntn1*) are necessary for improved regenerative outcomes, including axonal infiltration, vascularization, and improved behavioral recovery. Proteomic analysis of the extracellular vesicles show that astrocytes enrich pro-reparative proteins in extracellular vesicles with only 30 proteins relating to inflammatory or complement pathways loaded out of a total of 1073 proteins. Further, we show that the use of a biomaterial scaffold is necessary for the improved regeneration observed from reactive astrocyte extracellular vesicles. These studies show that reactive astrocytes use extracellular vesicles enriched with pro-repair proteins to promote recovery after injury.

## Main

Astrocytes are known to regulate the body responses to disease and injury in the central nervous system (CNS), as they become reactive and undergo functional changes.^1^ Reactive astrocytes isolated from lipopolysaccharide (LPS)-treated brain, which induces inflammation, display a neurotoxic profile, while reactive astrocytes isolated after ischemic stroke show a reparative profile.^2^ This data agrees with findings that the chemoablation of glial fibrillary acidic protein (GFAP) reactive astrocytes disrupts vascular repair and remodeling after ischemic stroke.^3^ Thus astrocytes hold promise as therapeutic sources to repair the brain after ischemic stroke. However, currently repair after stroke is limited to the region of the brain that contains reactive astrocytes, namely the glial scar. Notably, transplantation of patient-derived astrocyte progenitor cells after white matter stroke promotes endogenous remyelination and axonal sprouting by maturing into a pro-repair phenotype, which had better therapeutic outcomes than neural progenitor cells.^4^

Extracellular vesicles (EVs) are ubiquitously released by cells and are lipid bilayer nanoparticles containing proteins, RNAs, and metabolites, including cellular mediators that orchestrate downstream biological pathways. EVs have shown promise to promote repair and regeneration in a variety of diseases^5–7^ and several are in clinical trials.^8^ EVs released by astrocytes enhance neuronal survival and electrophysiological function.^9,10^ The activation of astrocytes with IL-1β and TNFα produce EVs that reduce dendritic growth of targeted neurons.^11^ Thus, the phenotype of parent cells dictate the EV cargo and biological function. Here we use soluble activation of primary astrocytes extracted from rat pup cortices to produce astrocytes with distinct gene expression profiles, collected their EVs, and studied their ability to promote repair after stroke in both peri-infarct and infarct areas.

*In vitro* the activation of astrocytes with interleukin-1 alpha (IL-1α), tumor necrosis factor alpha (TNF-α), and the classical complement component C1q^12^ show similar transcriptome profiles astrocytes activated with LPS *in vivo*. We choose this published cocktail to obtain the established IL-1α/TNF-α/C1q) induced astrocytes. While astrocytes isolated from middle cerebral artery occlusion (MCAO) stroke have shown pro-reparative gene profile^2^, there are no published reports of an *in vitro* reparative astrocyte cocktail. We used IL-4 and C1q to generate reactive astrocytes with a potentially distinct phenotype compared to the IL-1α/TNF-α/C1q induced astrocytes using the logic that C1q has induced up-regulation of many MCAO-specific genes in combination with IL-1β^12^, macrophages activate astrocytes^12^ and that reparative macrophages that promote tissue repair secrete IL-4.^13^

To verify the genomic phenotypes of the untreated (UT), IL-1α/TNF-α/C1q, and IL-4/C1q induced astrocytes, we first investigated their transcriptome by RNA sequencing (Fig. 1a and Extended Data Fig. 1). 13,176 genes were detected from these samples and 92.36% of them overlapped among the three astrocytes. IL-1α/TNF-α/C1q induced astrocytes showed significant up-regulation of *Lcn2, Cp*, and *Steap4*, indicating they became strongly reactive, and they had up-regulation of some ECM genes (*Col6a2, Fn1*), several matrix metalloproteinases, proteins involved in cell adhesion (*CD44, Nfasc*), genes related to immune responses (*Osmr, Cxcl10, Il6*) and complement pathways (*C3, Serping1*). Our IL-1α/TNF-α/C1q induced astrocytes had some different expression profile compared to previously reported neurotoxic A1 astrocytes (Extended Data Fig. 1e),^12^ such as downregulation of some pan-reactive markers (*Vim, Gfap*), fatty acid related markers (*Apoe, Scd2*),and LPS-specific markers (*Ggta1, Fbln5*). Also, there are several MACO-specific genes up-regulated in our IL-1α/TNF-α/C1q induced astrocytes (*Clcf1, Tgm1, Ptx3, Sphk1, Ptgs2, Cd14*). In contrast, IL-4/C1q induced astrocytes up-regulated pan-reactive markers (*Hspb1, Aspg, Vim, Gfap*) and some MACO-specific genes (*S100a10, Cd109, Emp1, Tm4sf1, B3gnt5*). They also had notable up-regulation of some ECM proteins, such as *Col12a1, Ecm1, Vcan, Ncan, Bcan*. Together, the transcriptome profiles of IL-1α/TNF-α/C1q, and IL-4/C1q activated UT astrocytes were distinct, allowing us to investigate the idea that reactive astrocytes release EV that promote recovery after stroke. Since our IL-1α/TNF-α/C1q induced astrocytes that had many reactive and inflammatory genes significantly up-regulated, we continue to term the astrocytes produced from this cocktail neurotoxic (NTx) astrocytes following previous studies^12,14^. IL-4/C1q induced astrocytes had fewer reactive genes up-regulated and showed strong up-regulation in many pro-reparative genes; thus, we term them reparative (Rep) astrocytes.

**Figure 1.**
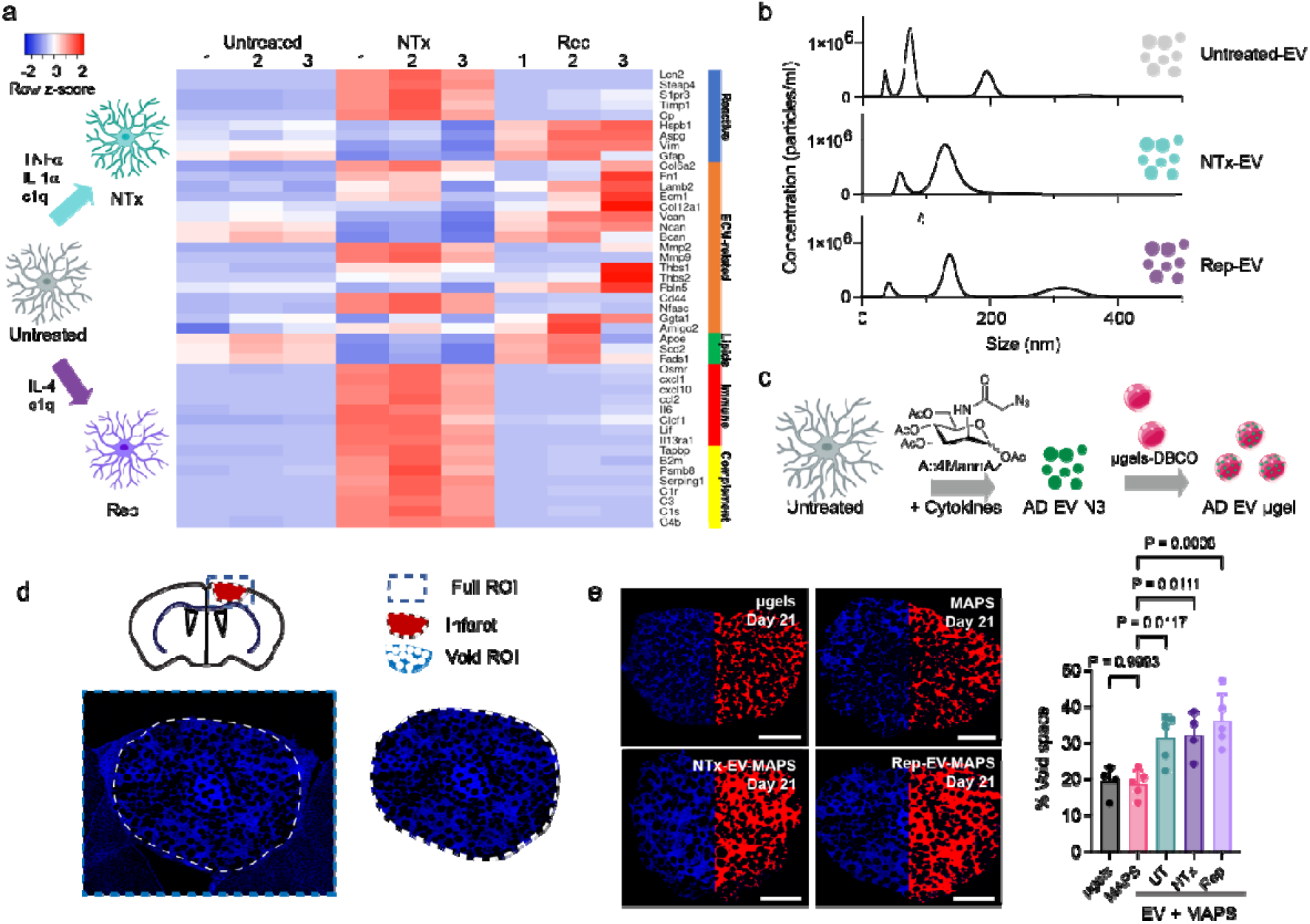
UT, NTx- and Rep-EVs promote matrix remodeling after hydrogels being injected into stroke infarcts. a) Schematic illustrating the generation of NTx and Rep astrocytes from naïve primary astrocytes, and the use of their derived EVs to make EV-MAPS for stroke treatment. Their phenotypes were distinguished from the RNA sequencing data generated heatmap showing the differential expression of selected genes in UT, NTx, and Rep astrocytes. b) Size characterization and concentrations of EVs derived from UT, NTx, and Rep astrocytes from nanoparticle tracking analysis. c) Schematic illustrating the metabolic labeling process to produce azide-bearing astrocytic EVs and the conjugation between astrocytic EVs and strain-alkyne microgels. d) Selection of the entire stroke infarct as region of interest for void space quantification to determine the degradation of MAPS. e) Z-projection confocal images and quantification suggesting the presence of astrocytic EVs on MAP gels significantly improved their degradation after implantation (day 5) into stroke cavities (n=4 or 5 mice). Scale bars, 500 μm. Statistics were calculated by one-way ANOVA with Tukey’s multiple comparisons test in **e**.

EVs collected from NTx, Rep, and UT astrocytes showed multiple size vesicles with the major size peaks between 50-150 nm, the exosome size range (Fig. 1b). The total astrocyte derived (AD) EV concentration was the same (10^7^ particles/mL) for all treatment conditions, indicating the cytokine treatment did not influence astrocyte derived EV secretion. We utilize a photo-thrombotic-stroke model located in the motor cortex as a reproducible model of ischemic stroke.^15^ The stroke in these studies spanned all cortex layers and resulted in a significant motor deficit in the left limb of the mouse (Extended Data Fig. 2a, b).

We first investigated injection of Rep-EV (1.8 × 10^6^ particles/mL) directly into the stroke cavity following our prior therapeutic approach^16,17^ to assess their therapeutic potential. In our therapeutic strategy, therapeutic agents are injected into the infarct at Day 5 post-stroke, a timepoint at which a cavity has formed and pro-repair pathways remain active. Rep-EV injection (16-days post injection) yielded similar outcomes in the infarct as the sham, no treatment group (Extended Data Fig. 2c-e). Rep-EV injection resulted a slight improvement of axon density in the peri-infarct region (Extended Data Fig. 2f-h), which was similar results to previous study.^18^ Though, repeated injections, higher dose, or a different timepoint, could improve outcomes by direct EV delivery, we moved to combine the EVs with a defined matrix that can be injected into the stroke core to deliver therapeutic cargo^19,20^ and that was previously found to reduce the inflammatory phenotype of astrocytes and microglia.^17^

Our defined matrix is a hydrogel composed of hyaluronic acid microgel building blocks, which undergoes gelation in situ through a bio-orthogonal strain-promoted azide-alkyne click reaction to form a porous hydrogel that allows for diffusion of soluble factors and cell infiltration. These hydrogels are termed microporous annealed particle scaffolds (MAPS).^21–23^ MAPS are fundamentally different than others used in the field because they are granular materials, injectable and porous, rather than continuous polymeric networks that must degrade for cellular remodeling to take place. Our hydrogels are modified with fibronectin derived RGD (Arg-Gly-Asp) peptide because we aim to create a chemically defined environment. Here strain-alkyne bearing microgels with an average size of 73.31 μm were used, which generated MAPS with 28% void space after *in vitro* assembled by multi-arm azide crosslinker. (Extended Data Fig. 3a, b)

Our goal was to retain reactive astrocyte derived EVs in our defined MAPS matrix and thus encourage repair beyond what is observed in the peri-infarct region. Thus, we covalently immobilized astrocytic EVs to MAPS via a strain-promoted azide-alkyne click chemistry. Astrocytic EVs were metabolically labeled with the azido-sugar, N-azidoacetylmannosamine-tetraacylated (Ac4MannAz), which has been previously shown to introduce azide groups at the glycocalyx of the EVs.^24,25^ Conjugation specific, only astrocytic EVs that were metabolically labeled with azides resulted in conjugation to strain-alkyne containing matrix (Extended Data Fig. 3c) and was effective, virtually no EVs were observed in solution after conjugation (Extended Data Fig. 3d, e). The amount loaded was dose dependent (Extended Data Fig. 3f) and impacted the gelation kinetics and final bulk storage modulus of our hydrogel matrix. As more astrocytic EVs are conjugated to the microgels, the modulus decreased due to reduced microparticle interlinking sites that were now occupied by EVs (Extended Data Fig. 3g). We chose 4×10^8^ EVs per milliliter of microgels for all experimental conditions as it allows for a significant number of EVs immobilization, while keeping the bulk modulus near that of the brain (300Pa). This EV dose was also consistent with the EV only treatment in Extended Data Fig. 2c.

**Figure 2.**
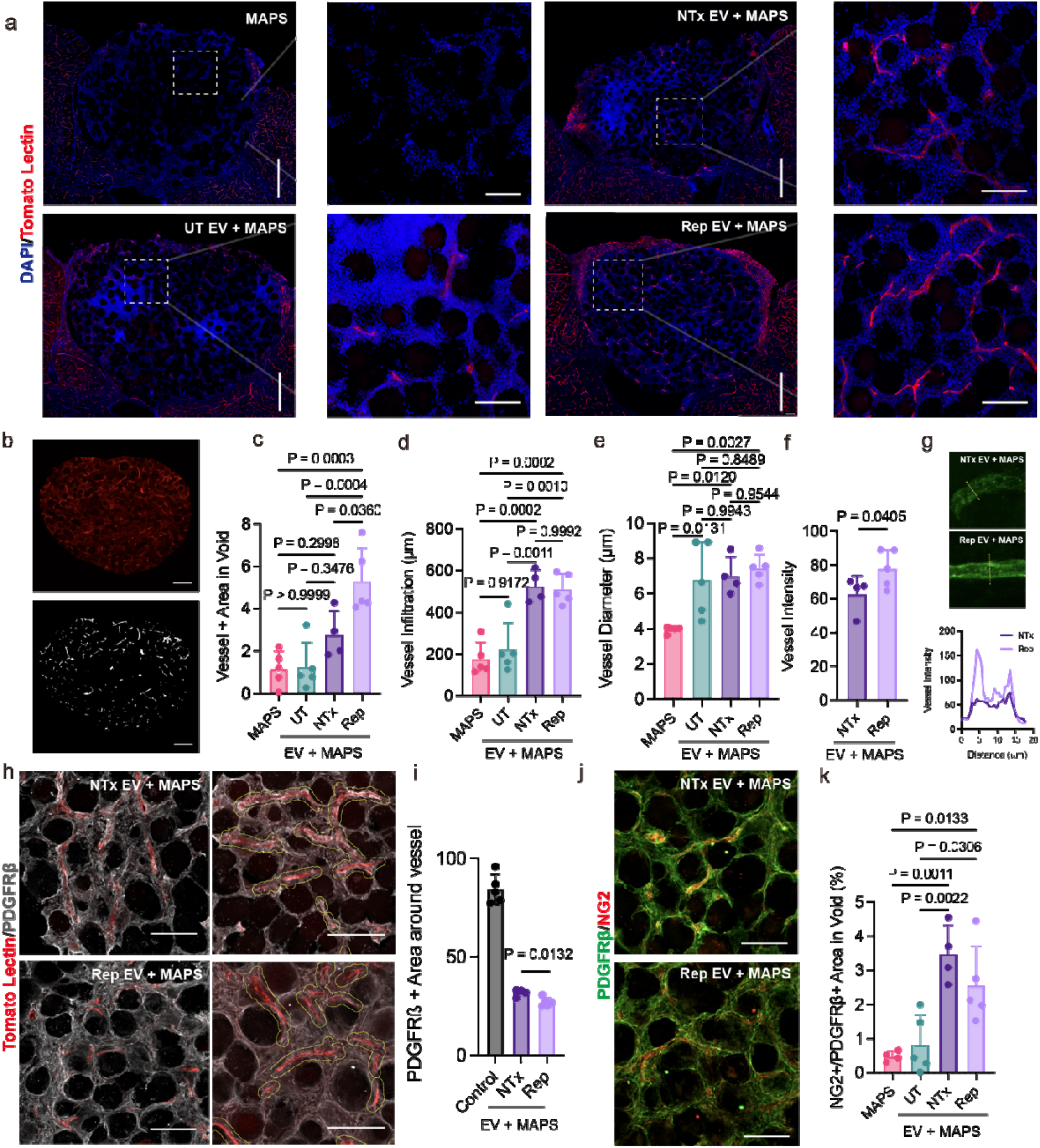
Rep-EV+MAPS improves angiogenesis and induces pericyte infiltration in infarct at Day 21 after stroke. a) Stitched 20X Z-projection large scan and inset images of perfused tomato lectin (red) in the stroke infarcts. Scale bars (large scans), 500 μm; Scale bars (insets), 100 μm. b) The tomato lectin signals within stroke infarct were binarized for blood vessel quantification. Scale bars, 250 μm. Quantification of c) total blood vessel area normalized to tissue area, d) blood vessel infiltration, and e) diameter of blood vessels showing improved angiogenesis in both NTx- and Rep-EV + MAPS groups (n=4 or 5 mice). f, g) Images and quantification of blood vessel intensity showing Rep-EV + MAPS had more mature vessels than NTx-EV + MAPS. h-i) Z-projection images and quantification showing pericyte (white) coverage over the new blood vessels (red) in the stroke infarct, indicating vessels are more mature in Rep-EV + MAPS group. (n=4 or 5 mice). Yellow lines are 5 μm boundary around the vessels as an illustration of vessel ROI for quantification of the interactions between pericytes and vessels. Scale bars, 100 μm. j-k) Z-projection images and quantification of pericytes by colocalizating PDGFRβ (green) and NG2 (red). Scale bars, 100 μm. Statistics were calculated by one-way ANOVA with Tukey’s multiple comparisons test in **c, d, e, f**; by one-tailed unpaired t-test between NTx-EV (n=4 mice) and Rep-EV (n=5 mice) groups in **k**; and by two-tailed unpaired t-test in **i**.

### Astrocytic EVs enhance MAP scaffolds degradation in stroke infarct

To evaluate the therapeutic outcomes of astrocytic EV + MAP on post stroke repair, we injected astrocytic EV modified microgels (4.5 μL injection) along with a tetra-polyethylene glycol-azide crosslinker into the cavity in a mouse photothrombotic stroke model at Day 5 post-stroke and collected the brain tissues at Day 21 post stroke (Extended Data Fig. 4a). MAPS scaffolds are infiltrated by stroke and peri-infarct resident cells through its void space, space between microgel particles, and not through direct microgel infiltration.^17^ Thus, the void space within the implanted MAPS is filled by residing and infiltrating cells (Fig. 1d and Extended Data Fig. 4b), we used the DAPI signal to estimate hydrogel remodeling by cells overtime. MAPS remodeling would result in more void which in turn results in more DAPI area. To account for the cell body and matrix deposition, we connected the DAPI signal using a Gaussian Blur to estimate the degree of MAPS remodeling in the stroke environment (Extended Data Fig. 4c). NTx- and Rep-EV + MAP scaffolds (NTx, and Rep) as well as UT EVs displayed improved hydrogel degradation compared to the MAPS only group (Fig. 1e). This improved degradation was not caused by the reduced crosslinking linkages between microgels as results of astrocytic EV conjugation, because a microgel only (no interlinking) group did not have significant degradation. These results show that the AD-EV delivery results in improved matrix remodeling and cellular infiltration within the stroke infarct.

### NTx- and Rep-EV-MAP scaffolds both promote angiogenesis after stroke

We have previously shown that angiogenesis is an essential component of biomaterial driven brain repair and behavioral improvement after ischemic stroke.^16^ In this work we focused on quantifying the density and morphology of perfused vessels that were connected to the circulation system after treatment with astrocytic EV decorated MAPS. Perfused vessels were labeled with tomato lectin before fixative perfusion, sectioning, and immunostaining. Gross observation revealed that NTx- and Rep-EV + MAPS groups had obvious vessels in the middle of the infarct, while UT-EV + MAPS and MAPS only groups did not (Fig. 2a). As expected, the vessels grew surrounding the microgels that form MAPS rather than penetrating through them (Fig. 2a insets), indicating that the void structure in MAPS can be remodeled in situ to support vessel growth. To quantify the density of the vasculature in the voids we measured the positive area of tomato lectin stain as well as the number of vessels per mm in a binarized image (Fig. 2b). To assess the vessel infiltration distance, we measure the distance from the edge of the stroke to the furthest vessel found in the section. Finally, to assess vessel morphology we assessed vessel branching and vessel thickness. Overall, new blood vessel formation within stroke infarct in NTx- and Rep-EV + MAPS groups was significantly improved compared to UT-EVs + MAPS and MAPS only groups (Fig. 2c-e and Extended Data Fig. 5a-d).

Both NTx- and Rep-EV groups had similar branch counts and blood vessel infiltration, but the total blood vessel area within the infarct and maximum branch length were higher in Rep-EV group, indicating increased angiogenic cargo in Rep-EVs compared to NTx-EVs. In addition, the new vessels in Rep-EV group had higher tomato lectin fluorescent intensity than those in NTx-EV group, indicating Rep-EV-MAPS promoted the regeneration of blood vessels with a more mature glycocalyx^3^ (Fig. 2g). Notably there are perfused vessels spanning the center of the infarct in both NTx- and Rep-EV conditions, a tissue that never recovers from stroke. This indicates that reactive astrocytic EVs impart a significant reparative response after stroke, after only 16 days EV + MAPS injection.

To further study the integrity of new blood vessels we stained for PDGFRβ, a marker of pericytes. We found a robust PDGFRβ+ cell population in the infarct region (Extended Data Fig. 5e, f). To determine what the % PDGFRβ+ area is in mature vessels we measure the % PDGFRβ+ area in the contralateral side using a vessel ROI such that only PDGFRβ+ area in vessels was captured. We found that 84% PDGFRβ+ area in the vessel ROI for the contralateral vessels (Fig. 2h, i and Extended Data Fig. 5g), while for Rep-EV and NTx-EV + MAPS treatments only 25-30% PDGFRβ+ area was found in the vessel ROI, indicating that the new blood vessels in the infarct are less mature than those in the contralateral side. Since our PDGFRβ stain appeared not to be specific to pericytes, we moved to define pericytes as cells co-expressing PDGFRβ and NG2.^26^ Co-expressing cells (PDGFRβ+ and NG2+) were found to colocalize with vessels and show to have a statistically higher positive area in the Rep-EV and NTx-EV + MAPS groups (Fig. 2j, k and Extended Data Fig. 5h, i). These results demonstrate that 16 days-post AD-EV + MAP injection, vessels are beginning to be remodeled to become mature.

The overall PDGFRβ+ area in the void showed a significant increase for Rep-EV and NTx-EV + MAPS condition compared to MAPS and UT-EV + MAPS condition, with 45% and 20% PDGFRβ+ area, respectively (Extended Data Fig. 5j). These results demonstrate that a PDGFRβ + cell population contributes significantly to the remodeled void space that leads to regenerative processes after stroke in the Rep-EV and NTx-EV + MAPS groups, which could possibly be oligodentrocyte progenitor cells and fibroblast.^27^ The majority of infiltrating cells in MAPS only group were Iba-1+ microglia/macrophages (Extended Data Fig. 6a), whereas EV + MAPS groups had limited amount of Iba-1+ cells, suggesting AD-EVs promoted transition from immune responses to regenerative process. The scar thickness was unchanged between EV + MAPS groups and MAPS only group, but all these groups had reduced scar compared to sham or EV only group (Extended Data Fig. 2c, d and 6b) as previously reported^17^.

### EVs derived from Neurotoxic Astrocytes improve axonal sprouting after stroke

Neurotoxic astrocytes were found to kill neurons by secreting APOE and APOJ saturated lipids^14^ and post stroke axonal sprouting is necessary for behavioral improvement after stroke.^28^ Thus, whether NTx- and Rep-EVs promoted post stroke axonal sprouting was of interest. Both NTx- and Rep-EV groups resulted in increased neurofilament area in the infarct and peri-infarct compared to MAPS only (Fig. 3a-e). However, NTx-EV + MAPS group had the highest NF200 neurofilament area in infarct and peri-infarct as well as NF200 infiltration distance (Fig. 3c-e), despite being derived from neurotoxic astrocytes suggesting that harmful saturated lipids are not presented in NTx-EVs.

**Figure 3.**
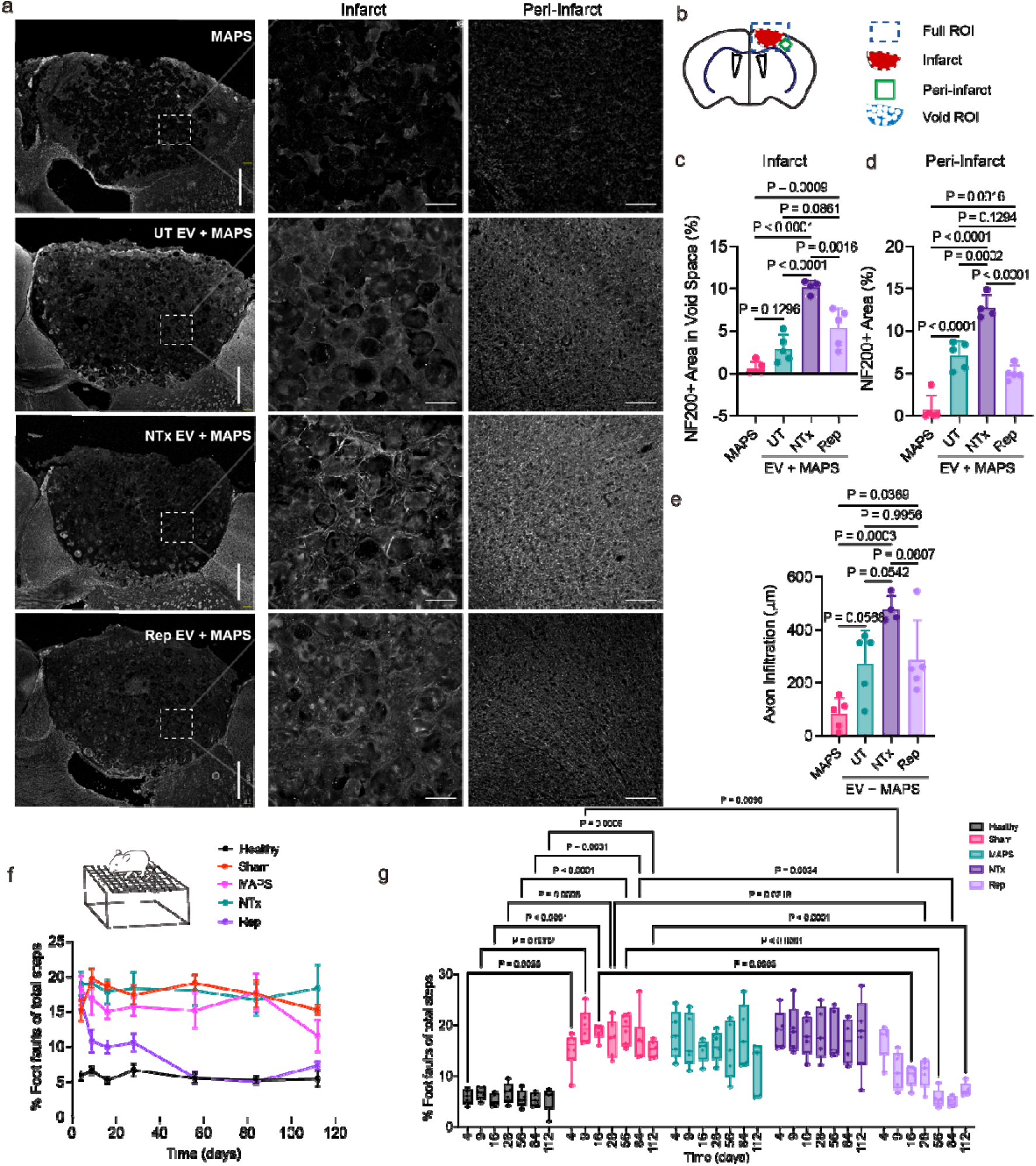
NTx-EV + MAPS improves axon regrowth at Day 21 after stroke while Rep-EV + MAPS promotes functional recovery. a) Stitched 20X Z-projection large scan and inset images of NF200 (white) in the stroke and peri-infarct regions. Scale bars (large scans), 500 μm; Scale bars (insets), 100 μm. b) Schematic showing the region of interest for the stroke infarct, peri-infarct, and the tissue area between MAPS for image quantification. Quantification of c) total axon area in infarct normalized to tissue area, d) axon area in periinfarct, and e) axon infiltration showing NTx-EV + MAPS promotes axon regrowth in both infarct and peri-infarct after stroke (n=4 or 5 mice). f-g) Grid walking behavioral studies showing functional recoveries from the treatment of Rep-EV + MAPS after stroke (n= 5 mice). Statistics were calculated by one-way ANOVA with Tukey’s multiple comparisons test in **c, d, e**; and by two-way ANOVA with Dunnett’s multiple comparisons test in **g**.

### Rep-EV promote functional recovery after stroke

To evaluate the functional recovery after stroke as results of implanting ADEV-MAPS scaffolds, we performed grid walking behavioral studies on mice treated with each experimental condition and quantified the ratio of missed and total steps (Fig. 3f, g). We first created a baseline comparison between healthy and stroked mice, in which the stroked mice had around 2.5-fold higher missed/total step ratio compared to healthy mice throughout the 4-month period. The three EV + MAPS groups (UT, Rep, NTx) had similar percentage of missed steps at Day 4 compared to stroked mice, but Rep-EV + MAPS group significantly decreased the percentage of missed steps at Day 16 compared to sham, while MAPS and NTx-EV + MAPS did not show any improvement. The mice in Rep-EV + MAP group continued to recover and have similar missed steps as healthy mice after two months (Fig. 3f, g).

These results collectively showed that reactive astrocytes that express both inflammatory and reactive genes, have EVs with pro-reparative cargos capable of promoting tissue repair, while reactive astrocytes that express less inflammatory genes have EVs that promote functional recovery after stroke.

### Proteomic analysis on ADEVs

Due to the differential tissue repair results from EV + MAPS scaffolds derived from UT, NTx, and Rep astrocytes, we investigated the protein cargo differences to mechanistically explain the key protein components and pathways leading to regeneration in each group. We lysed the astrocytic EVs after collection and performed proteomic analysis by nanoscale liquid chromatography coupled to tandem mass spectrometry (nanoLC-MS/MS) and label-free quantification (Fig. 4a). 1073 proteins were detected in total from all samples and 94% of these proteins are congruent between Rep, NTx, and UT-EV groups (Extended Data Fig. 7a). The label-free quantification showed the protein abundances in NTx-EVs were drastically different compared to UT EVs, whereas the protein profile in Rep-EVs was similar to UT EVs (Fig. 4b and Extended Data Fig. 7b, c). These trends are similar to the gene expression data (Fig. 1a), indicating the astrocytic EVs contain protein cargos that correlate with their parent cells. Comparing proteins in Rep-EVs to NTx-EVs using significance ratio of 2.0 and fold change of 1.5, there were 4 up-regulated proteins, including Gas6 involved in reoxygenation^29^, PTPRS relating to synapse formation^30^, FGFBP1 involved in regeneration^31^, and SLIT2 involved in BBB repair^32^, and 11 down-regulated proteins, containing neuronal growth factor VGF (Fig. 4b, c). These proteins are possible key players contributing to the angiogenesis in Rep-EV + MAPS (Fig. 2) and axongenesis in NTx-EV +

**Figure 4.**
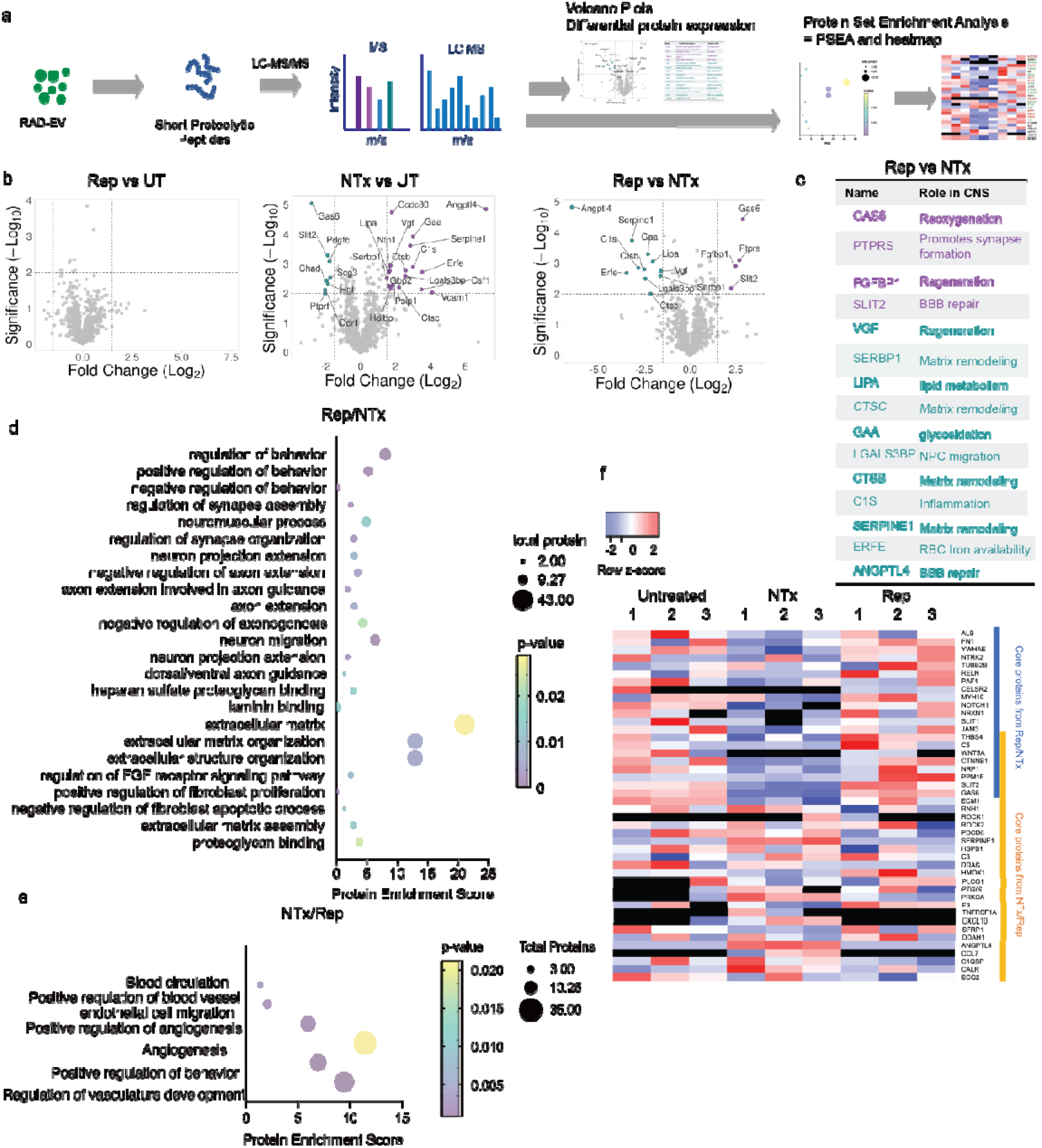
Proteomics profiling of proteins components within UT, NTx, and Rep-EVs. a) Schematic showing the process of sample preparation, data acquisition on LC-MS/MS, and differential and enrichment analysis. b) Volcano plots showing differential expression of proteins between UT, NTx, and Rep-EVs. c) The roles in CNS of up- and down-regulated proteins comparing Rep-EVs and NTx-EVs. PSEA-Quant analysis of up-regulated pathways using abundance ratios of d) Rep/NTx and e) NTx/Rep. f) Heatmap of core proteins from PSEA-Quant analysis showing the differential expression levels of these key proteins among UT, NTx- and Rep-EVs.

MAPS (Fig. 3) treated animals. Interestingly, many inflammatory genes found in RNA sequencing data were not presented as proteins in EVs (Extended Data Fig. 7d), especially for NTx astrocytes. NTx astrocytes had a 9.005 log2 fold change of C3, the most characteristic and highly up-regulated marker for NTx astrocytes^12^, at the cell level compared to UT astrocytes, but only 0.396 log2 fold change in EV level (Extended Data Fig. 7e), suggesting the reduced toxicity in NTx-EVs and explaining why axon regeneration was induced by NTx-EV + MAPS.

Because most of the proteins are the same among UT, NTx, and Rep-EVs, we analyzed the abundance ratios for enrichment analysis to obtain significantly up-regulated pathways in a group. Therefore, we used protein set enrichment analysis (PSEA)-Quant of abundance ratios between Rep/NTx and NTx/Rep to find gene ontology pathways of reparative processes (Fig. 4d, e). Surprisingly, Rep-EVs have 14 neuronal growth pathways significantly up-regulated, including axon extension involved in axon guidance (GO:0048846), regulation of synapse assembly (GO:0051963), dorsal/ventral axon guidance (GO:0033563), neuron projection extension (GO:1990138), neuron migration (GO:0001764), and axon extension (GO:0048675). Furthermore, there were 7 ECM protein related pathways up-regulated in Rep-EVs, including extracellular matrix (GO:0031012), proteoglycan binding (GO:0043394), extracellular matrix organization (GO:0030198), extracellular structure organization (GO:0043062), and heparan sulfate proteoglycan binding (GO:0043395). Rep-EV + MAPS group was the only treatment group that had a significant behavioral improvement, suggesting that these reparative Gene Ontology pathways play a significant role in recovery after stroke. 1

NTx-EVs also up-regulate pathways compared to Rep-EVs, which were mostly angiogenesis related, including angiogenesis (GO:0001525), positive regulation of behavior (GO:0048520), and regulation of vasculature development (GO:1901342), which may contribute to the improved PDGFRβ+ cells surrounding vessels in NTx-EV + MAPS group.

Core proteins from PSEA-Quant were plotted in a heatmap to show the difference of expression levels among the three groups (Fig. 4f). Rep-EVs group had higher expression of most of the core proteins from Rep/NTx analysis including some regenerative proteins, such as FN1, THBS4, and RELN, indicating these proteins probably contributed to the tissue regeneration and behavioral recovery in Rep-EVs group. Apart from these core proteins, NTx-EVs had more expression of axon permissive factors NTN1^33^ and VGF and less expression of axon inhibitory factors, such as BCAN, VCAN, SLIT2^34^, which possibly leads to improved axon infiltration in NTx-EVs group (Fig. 3).

## Discussion

Here we show Rep-EVs drive substantial regeneration of blood vessels and axons in the infarct after stroke and result in functional behavior recovery. The regeneration only occurs when Rep-EVs are immobilized onto a biomaterial scaffold, which is in line with other research showing materials prolong EV retention *in vivo*^35,36^. Our MAPS preserves the injectability of EVs after immobilization and the unique inner microporous structure present the infiltrating cells an environment of interconnected channels coated with astrocytic EVs. The presence of astrocytic EVs enhances matrix remodeling by the infiltrating cells and transits the infarct from an inflammatory to a reparative environment as the major population of infiltrating microglia decreases significantly. Astrocytic EVs also enrich pro-reparative proteins that are related to signaling pathways for tissue regeneration and functional recovery. However, why astrocytes choose to pack pro-reparative proteins instead of reactive and pro-inflammatory proteins remain unclear. NTx astrocytes upregulate reactive and complement genes in cell level, but most of these markers are low or not presented in NTx-EVs.

Our findings highlight a versatile injectable biomaterial platform to deliver EVs from all cell types for tissue repair. Future studies can try other cytokine combinations for astrocyte activation and use EV + MAPS method to test their functions in CNS diseases. This could be an easier way to screen astrocyte functions at different states. EVs from other cell types, such as pericytes, microglia, oligodendrocytes, and T cells, can also be metabolically labeled and attached to MAPS to study their therapeutic outcomes for treating CNS diseases.

## Methods

### Primary astrocyte extraction and culture

Rat cortical astrocytes were prepared from P1 rat cortices as previously reported^37^. Briefly, the cortices were micro-dissected and papain digested followed by trituration in low and high ovomucoid solutions. Cells were passed through a 20 μm mesh filter, resuspended in astrocyte growth media (AGM; DMEM, 10% FBS, 10 μ *M* hydrocortisone, 100 U/mL Pen/Strep, 2 *mM* L-glutamine, 5 μg/ml insulin, 1 *mM* NaPyruvate, 5 μg/ml N-acetyl-L-cysteine) and 15-20 million cells were plated on 75 mm^2^ flasks (non-ventilated cap) coated with poly-D-lysine. Flasks containing cells were incubated at 37 °C in 5% CO_2_. On DIV3, AGM was removed and replaced with DPBS. Flasks were then shaken by hand for 10-15 times until only the adherent monolayer of astroglia remained. DPBS was then replaced with fresh AGM. Cytosine arabinoside was supplemented to the media on DIV5 for two days to eliminate fast dividing cells.

### ADEV isolation from primary astrocyte culture media

On DIV6, AGM were replaced by EV-collecting media (DMEM, 10% exosome-depleted FBS, 10 μ*M* hydrocortisone, 100 U/mL Pen/Strep, 2 mM L-glutamine, 5 μg/ml insulin, 1 m*M* NaPyruvate, 5 μg/ml N-acetyl-L-cysteine, 50 μM Ac4ManNAz) with the inducing cytokines (3□ng/ml IL-1α, 30□ng/ml TNF, and 400□ng/ml C1q for NTx activation, 20 ng/mL IL-4 and 400 ng. ml C1q for Rep activation. After 24 hrs, the culture media were collected and centrifuged at 2000g for 30 mins to remove cells and debris. The supernatant was transferred into a new tube and mixed with half volume of the total exosome isolation kit (Thermo Scientific) for overnight incubation at 4 °C. The solutions were then centrifuged at 10,000g for 60 mins at 4 °C and the resulted ADEV pellets were ready to be resuspended in PBS for downstream process.

### Preparation of ADEV-MAP hydrogels

The microfluidic device was made of polydimethylsiloxane (PDMS) by standard soft lithography. Master molds were fabricated on a 4-inch silicon wafer by a photolithographic technique using a negative photoresistor (SU8 2075). Microfluidic devices were molded from master molds by pouring degassed PDMS (elastomer:crosslinker = 10:1) and cured at 60 °C for 1h. PDMS devices were then placed onto a glass slide and bonded together at 60 °C overnight. The bonded device was treated with Rain-X to render a hydrophobic surface. Microgel droplets were generated at the flow focusing region where the oil phase broke off the gel solution into droplets. The gel solutions consisted of 3.4 wt % hyaluronic acid-norbornene (MW = 79 kDa, degree of norbornene functionality was 35%), MMP-degradable crosslinker GCRDGPQGIWGQDRCG (2.3 *mM*), lithium phenyl-2,4,6-trimethylbenzoylphosphinate (10 m*M*), and RGDSPGERCG (1 m*M*). The oil phase included heavy mineral oil with Span 80 (15 wt %). The flow rate for gel precursors was 0.3 μL/min and that for oil phase was 2.7 uL/min. The microgel droplets were cured downstream with ultraviolet (UV) irradiation (20 mW/cm^2^, 365 nm). The prepared norbornene-bearing microgels were washed and sterilized with PBS, 1% pluronic F127, and 30% ethanol. DBCO-tetrazine was used to convert the free norbornene groups to DBCO groups through Diels-Alder click reaction at 37 °C for 4 hours. DBCO-excessive microgels were incubated with azido-ADEV suspension at 37 °C for 30 mins to make ADEV-conjugated microgels. These microgels were crosslinked with 4-arm PEG-azide to assemble into MAP hydrogels.

### Characterization of ADEVs and ADEV-hydrogels

The quantity and size of ADEVs were characterized using nanoparticle tracking analysis. ADEVs were fluorescently stained with plasma membrane dye to visualize their conjugation on microgels by confocal imaging. *In situ* monitoring of ADEV-MAP hydrogel gelation was performed on rheometer at 1% strain and 2.5 rads/s frequency.

### Mouse photothrombotic stroke model

PT stroke was induced using an intraperitoneal (i.p.)-injected light-sensitive dye. Briefly, mice were positioned in a stereotaxic instrument and administered Rose Bengal (10□mg/ml, 10 uL. g i.p.). After 7 minutes, a laser was illuminated on the spot of the closed skull 1.5□mm left from the bregma for 13 minutes.

### Transplantation of MAPS after stroke

At Day 5 post-stroke surgery, 4.5□μl of MAPS that conjugated with UT, NTx, or Rep-EVs were injected directly into the stroke cavity through a 30-gauge needle attached to a 10□μl Hamilton syringe at 1□μl/min. The needle was positioned at 0.75 mm penetration from the skull. The control groups were injected with PBS or the same amount of ADEVs in PBS solutions.

### Tissue processing and immunohistochemistry

At Day 21 post-stroke, mice were retro-orbitally injected with biotinylated tomato lectin (0.1 mg) 10 mins before perfusion and then transcardially perfused with cold PBS and 4% paraformaldehyde. Brains were harvested and postfixed in 4% PFA for 2 hrs and immersed in 30% sucrose until sinking for cryoprotection. Tangential sectioning of frozen brains (30 μm thick) was obtained using a cryostat and mounted on gelatin-coated glass slides. Immunohistochemistry included antigen-retrieval by incubating the tissue sections in citrate buffer pH = 6 for 30 mins at 80 °C, washing steps using PBS, and permeabilization/blockage using Triton (0.3%) and normal donkey serum (10%). Primary antibodies were as follows: rat anti-GFAP (1:400) for astrocytes; rabbit anti-Iba-1 (1:250) for microglias; rabbit anti-NF200 (1:200) for neurofilaments; goat anti-PDGFRβ (1:20) and rabbit anti-NG2 (1:500) for pericytes. Primary antibodies were incubated overnight at 4□°C followed by a fluorescently labelled secondary antibody (1:500) for 1□h at room temperature with streptavidin-conjugated dye (1:500) for biotinylated tomato lectin and DAPI (1:1000) for cell nuclei. After washing, Aquapoly was used as mounting media to mount coverslip onto the tissue sections.

### Confocal imaging and quantitative analysis

Large confocal scans on the entire stroke infarct were performed by stitching 4 * 5 20X field-view. Brain sections from 900-1200 μm into the stroke infarct were used for tissue study quantification. The DAPI signals of infiltrating cells within the ischemic core were used to quantify the void space of MAPS and EV + MAPS groups to determine the degradation of MAPS. By applying a gaussian blur to the DAPI signal, a mask for tissue within the infarct area was created. The variance (σ) of a gaussian blur determines how much blur there is. This value was chosen as the one which obscured individual cells without removing features of the void (σ = 8, Extended Data Fig. 4c). This mask was used to normalize area-based measurements made within the infarct. Angiogenesis and axonogenesis were quantified by using tomato lectin and NF200 signal, respectively. Positive area was measured by denoising using the ImageJ Despeckle filter, and thresholding the image. Infiltration into the infarct was determined by taking the shortest distance between the edge of the ROI and signal within it.

Blood vessels were further quantified by skeletonizing the tomato lectin signal and using the AnalyzeSkeleton plugin in ImageJ. To reduce noise, only vessels greater than 50 μm in length were considered. From the results, vessel average length, maximum length, number of branches, and count over area were calculated. For each sample, vessel thickness was the average of 20 random measurements of blood vessel cross-sections. The vessel intensity profile was also taken from cross-sections of blood vessels. PDGFRβ+ area surrounding the tomato lectin signal was measured by first selecting the vessels within a sample, enlarging the selection by 5 μm on each side, applying the selection onto the PDGFRβ image, and then thresholding appropriately. After thresholding NG2+ and PDGFRβ+ areas separately, the percent of the infarct positive for both stains was used for colocalization analysis.

### Proteomic analysis

The ADEV pellets obtained after isolation were resuspend in 8 *M* urea, 50 m*M* ammoniom bicarbonate in H2O with protease inhibitors. These solutions were mixed and sonicated for 5 cycles (30 sec on, 30 sec off). The total protein quantity within these samples were determined by micro BCA assay. The samples were then spiked with 6-8 *M* urea/50 m*M* Tris-HCl (pH = 8) and reduced with 5 m*M* dithiolthreitol for 30 mins at 37 °C and alkylated with 15 m*M* iodoacetamide for 30 mins at room temperature in the dark. After 3X urea and Tris washes, proteins were figested using trypsin for overnight at 37 °C and eluted using 0.001% Zwittergent 3-16 and 1% HCOOH. Samples were dissolved in 200 μL 0.001% Zwittergent 3-16 prior to nanoLC-MS/MS.

Quantitative LC–MS/MS was performed on 2 μl of each sample, using an EASY nanoLC-1200 coupled to a Thermo Scientific Orbitrap Exploris-480 via an EASY-Spray source. In brief, the sample was first trapped on a Thermo Scientific Acclaim PepMap 100 C_18_ nanoLC 20 mm × 75 μm trapping column (5 μl/min at 99.9/0.1 v/v water/acetonitrile), after which the analytical separation was performed using an EASY-Spray C_18_ nanoLC 75 μm × 250 mm column with a 150-min linear gradient of 5–95% acetonitrile with 0.1% formic acid at a flow rate of 300 nl/min with a column temperature of 55□°C. Data collection on the Orbitrap Exploris-480 mass spectrometer was performed in a data-dependent acquisition (DDA) mode of acquisition with r = 120,000 (at m/z 200) full MS scan from m/z 375 to 1,600 with a target AGC value of 2 × 10^5^ ions. MS/MS scans were acquired at rapid scan rate in the linear ion trap with an AGC target of 5 × 10^3^ ions and a max injection time of 100 ms. The total cycle time for MS and MS/MS scans was 1.5 s. A 20 s dynamic exclusion was employed to increase depth of coverage.

Proteome Discoverer 2.5 (Thermo Scientific) was used for database searching of the experimental nanoLC-MS/MS data against protein sequences for *Rattus norvegicus* (SwissProt, reviewed, TaxID = 10116). Mascot Distiller and Mascot Server were used to produce fragment ion spectra and to perform the database searches using full trypsin enzyme rules with 5 ppm precursor and 0.02 Da product ion match tolerances. Database search parameters included static modification on cysteine (carbamidomethyl) and dynamic modifications on N-terminal acetylation, methionine loss, and methionine loss plus acetylation. Peptide Validator and Protein FDR Validator nodes in Proteome Discoverer were used to annotate the data at a maximum 1% protein false discovery rate.

## Acknowledgements

We would like to thank the National Institutes of Health, the National Institutes of Neurological Disorders and Stroke (1R01NS112940, 1R01NS079691, R01NS094599), the National Institute of Allergy and Infectious Disease (1R01AI152568), and the AXC Grant (CEA383000297) for funding in the early stages of the project. We thank Dr. Hongxia Bai at Molecular Education, Technology and Research Innovation Center (METRIC) at NC State University for proteomics experiments and analysis. We thank Dr. Tony Jun Huang for the use of nanoparticle tracking analysis.

## Author Contributions

S.X. and T.S. conceived the work. S.X. performed most experiments and N.V.P. and L.Z. helped with experiments. L.Z. performed data analyses. S.X. and T.S. wrote the manuscript, S.T.C. critically revised the manuscript with input from all other authors.

---

**Extended Data Figure 1.**
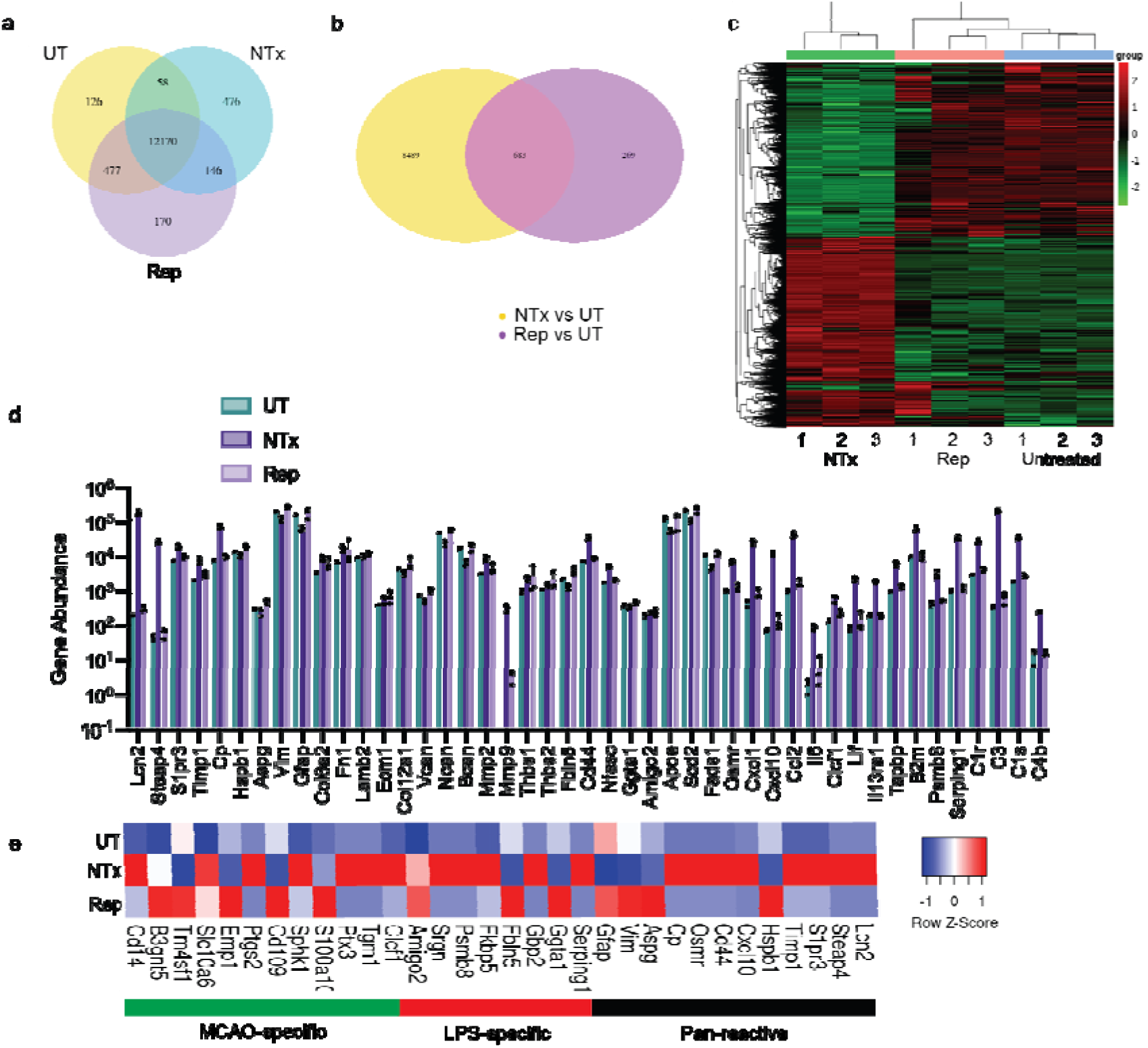
Differential gene profiles of UT, NTx, and Rep astrocytes from RNA sequencing. a) Venn diagram showing the overlapping genes among three astrocytes. b) Venn diagram showing comparing to UT astrocytes, NTx astrocytes had more up-regulated genes than Rep astrocytes. c) Heatmap showing the gene profile of NTx astrocytes was distinct from UT ones, whereas Rep astrocytes were more similar. d) The abundance comparison among three astrocytes of functional genes showed in Fig.1a heatmap. e) Heatmap showing the up-regulation of pan-reactive, LPS-specific, and MCAO-specific markers in our UT, NTx, and Rep astrocytes as a phenotype comparison to the astrocytes in Liddelow *et al*. paper.^12^

**Extended Data Figure 2.**
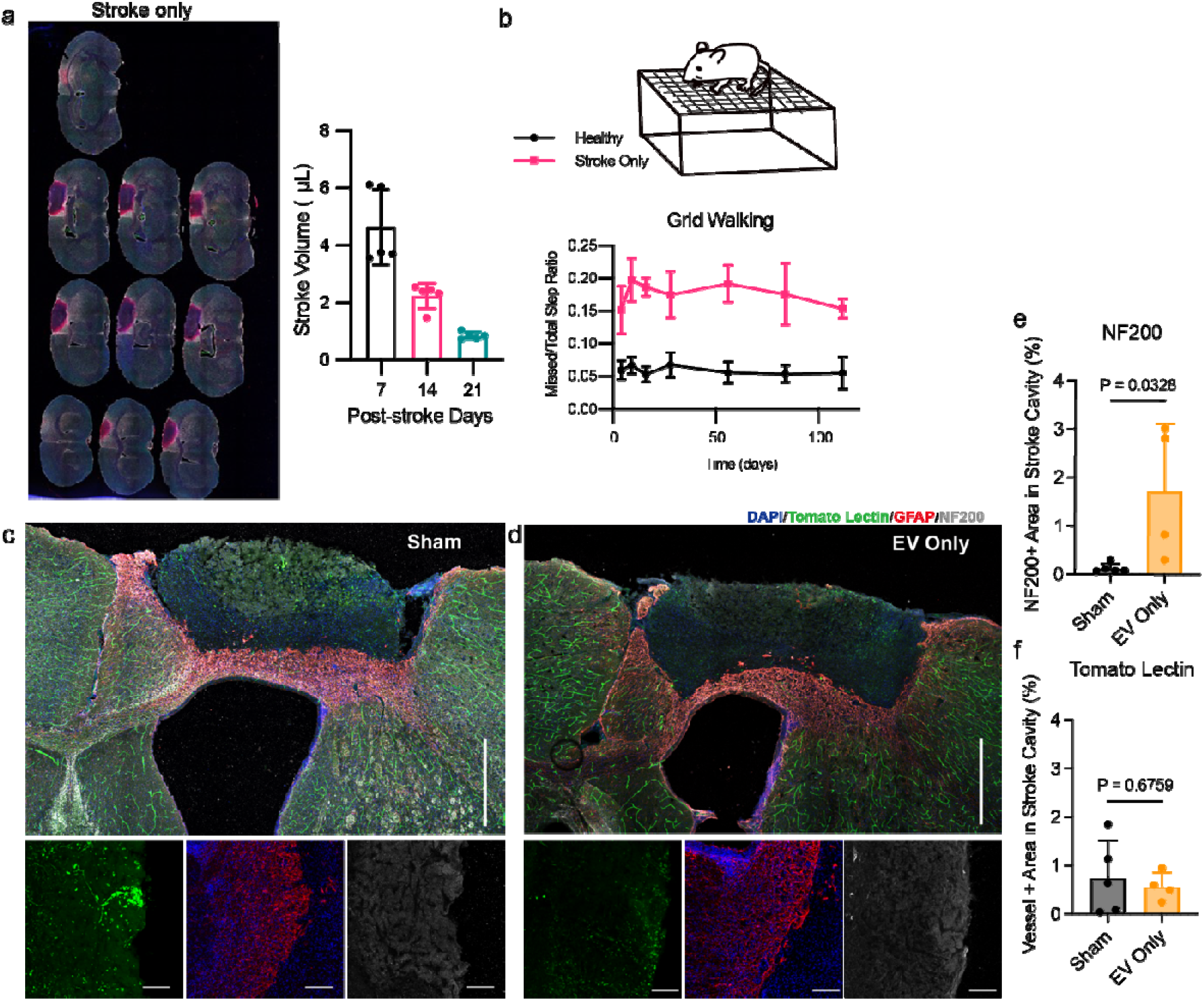
Injection of Rep-EVs only after stroke is not enough to repair blood vessels and axons. a) Stitched 4X Z-projection large scan image of a stroke only sample and quantification of stroke volume post-stroke. b) Schematic depicting the grid walking setup and quantification of healthy (n=5 mice) and stroke only (n=6 mice) showing functional differences after stroke. Stitched 20X Z-projection images of large image scans and inset images of tomato lectin (green), GFAP (red), and NF200 (white) in the stroke and peri-infarct regions for c) sham and d) EV only groups. Scale bars (large scans), 500 μm; Scale bars (insets), 100 μm. Quantification of e) total axon area, and f) total vessel area in the infarct between Sham (n=3 mice) and EV only (n=4 mice). Statistics were calculated by twotailed unpaired t-test in **e, f.**

**Extended Data Figure 3.**
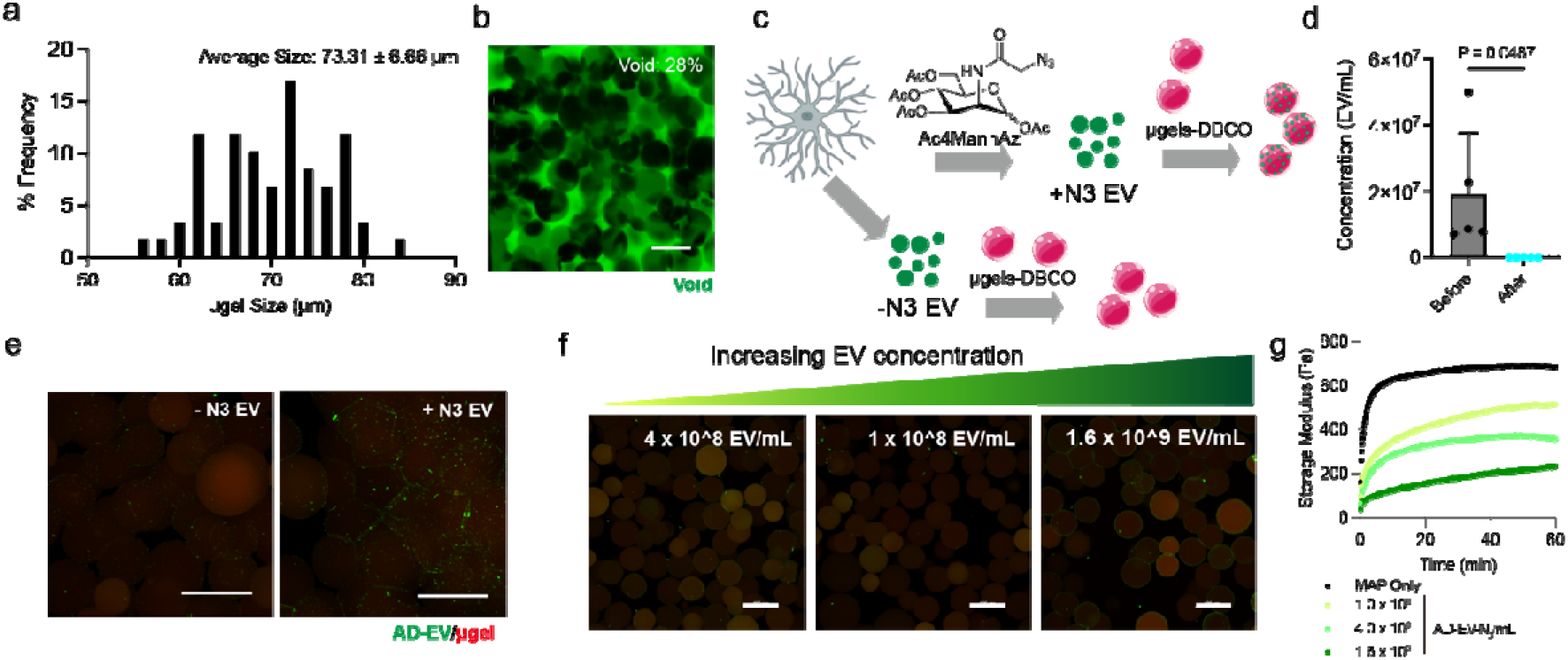
Conjugation of AD-EVs onto hyaluronic acid microgels. a) Histogram of microgel size. b) Confocal images of void space in MAPS infiltrated by high molecular weight dextran. c) Schematic illustrating the metabolic labeling process to produce azide-bearing astrocytic EVs and the conjugation between astrocytic EVs and strain-alkyne microgels. d, e) Z-projection confocal images and concentrations of EV suspensions indicating the successful conjugation of ADEVs onto microgels via strain-promoted azide-alkyne click chemistry. Scale bars, 100 μm. f) Z-projection confocal images showing the different thickness of EV layers at different EV concentrations on the surface of microgels. Scale bars, 100 μm. g) *In situ* rheological monitoring of the MAPS annealing process with the presence of different concentrations of EVs on the microgels. Statistics were calculated by two-tailed unpaired t-test in **d**.

**Extended Data Figure 4.**
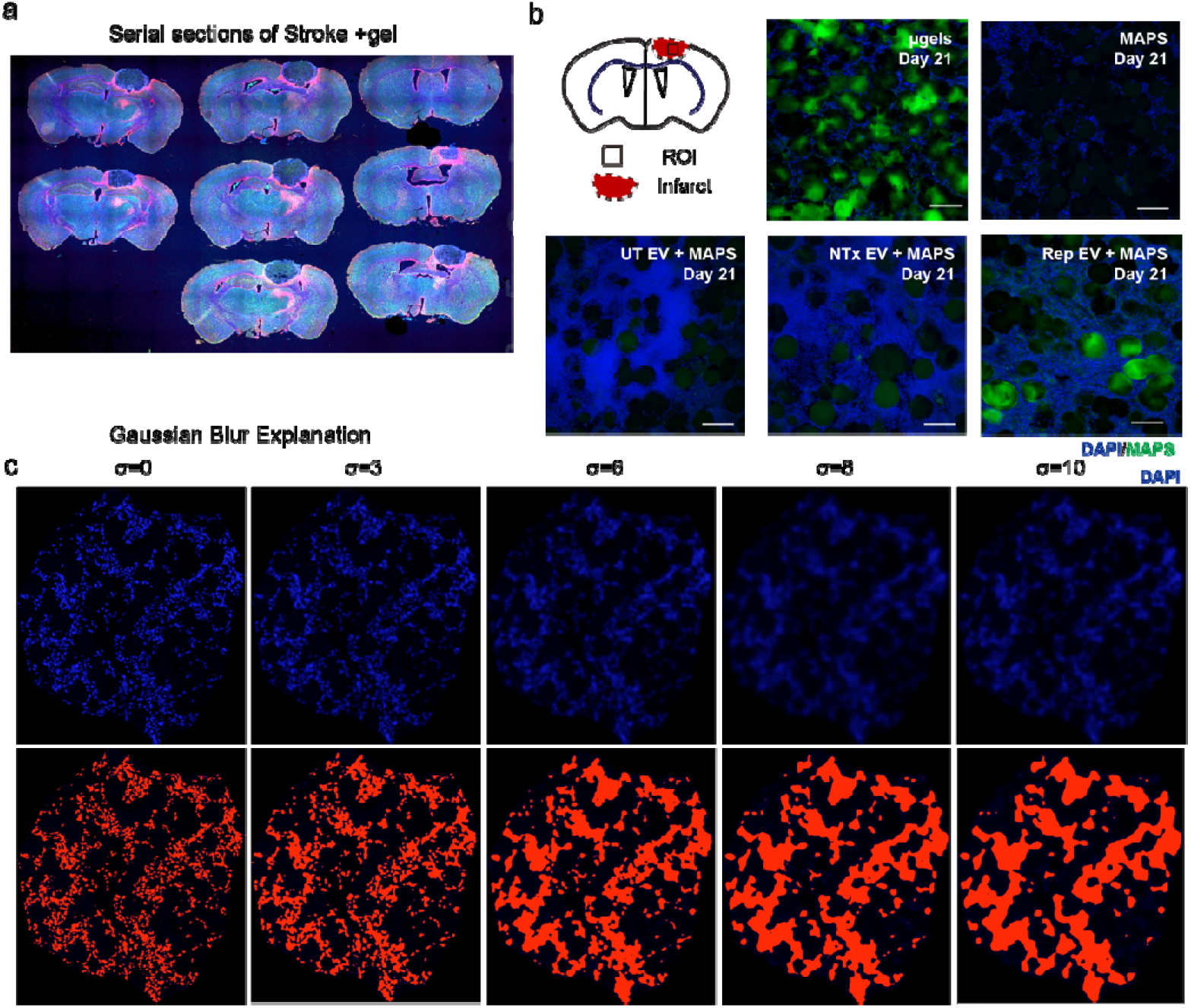
Zoomed in images and quantification method illustration of MAPS degradation after implanted into the stroke infarct. a) A stitched 4X Z-projection large scan image of a sample at Day 21 after stroke with MAPS injected into the infarct. b) Z-projection 20X confocal images showing the MAPS degradation after implantation. Scale bars, 100 μm. c) Z-projection large scan images of DAPI staining showing the gaussian blur and mask at different variances (σ).

**Extended Data Figure 5.**
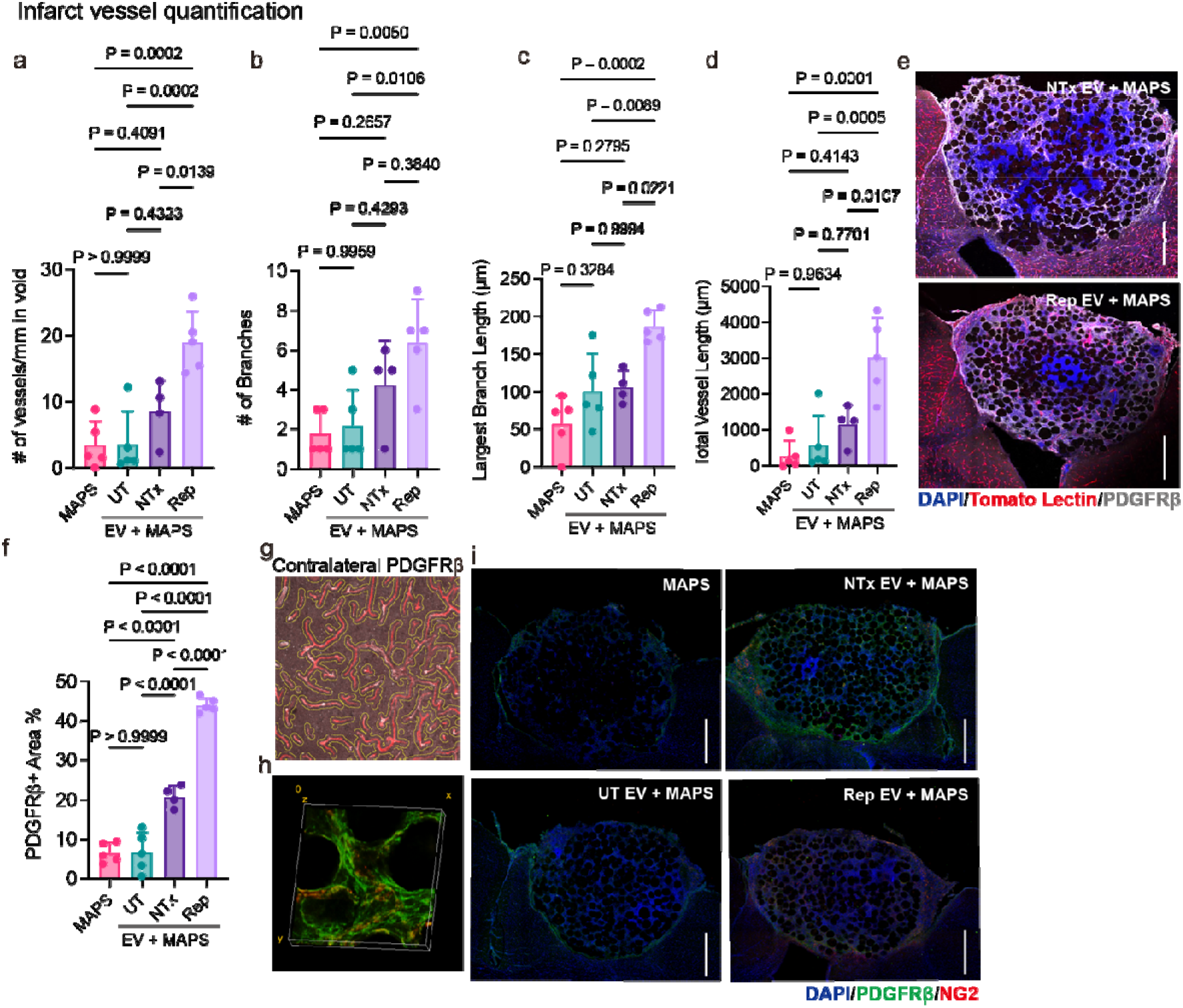
Rep-EV+MAPS induce neovascular network with more vessels, more branches, and longer length. Quantification of a) vessel count normalized to tissue area, b) branch count, c) length of longest vessel, and d) total vessel length showing improved angiogenesis in the Rep-EV + MAP group (n=4 or 5 mice). e-f) Z-projection images and quantification showing pericytes (white) constituted a large population of infiltrated cells and they were not restricted to the area around newly formed blood vessels (n=4 or 5 mice). Scale bars, 500 μm. g) Z-projection image showing pericytes were mostly restrict to the blood vessel surroundings in the peri-infarct. h) 3D rendering image showing the colocalization of PDGFRβ and NG2 signals. i) Stitched 20X Z-projection large scan images of pericyte staining in the stroke infarct using PDGFRβ (green) and NG2 (red). Scale bars, 500 μm. Statistics were calculated by one-way ANOVA with Tukey’s multiple comparisons test in **a, b, c, d**; by two-tailed unpaired t-test in **f.**

**Extended Data Figure 6.**
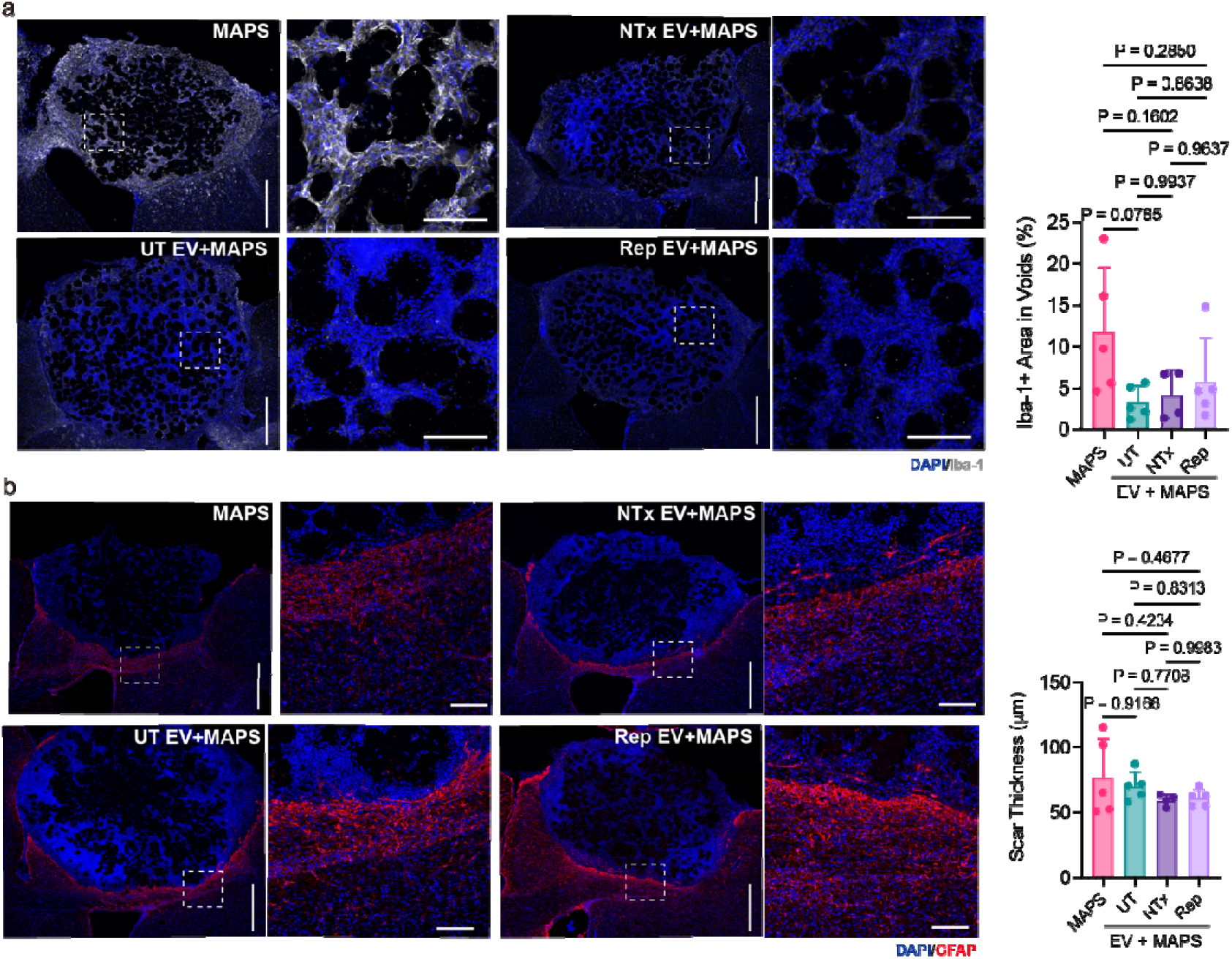
UT, NTx-, and Rep-EV+MAPS induces a less inflammatory environment in stroke infarct. a) Stitched 20X Z-projection large scan and inset images and quantification of Iba-1 (white) in the stroke infarct (n=4 or 5 mice). Scale bars (large scans), 500 μm; Scale bars (insets), 100 μm. b) Stitched 20X Z-projection large scan and inset images and quantification of GFAP (red) in the stroke infarct (n=4 or 5 mice). Scale bars (large scans), 500 μm; Scale bars (insets), 100 μm. Statistics were calculated by one-way ANOVA with Tukey’s multiple comparisons test in **a, b**.

**Extended Data Figure 7.**
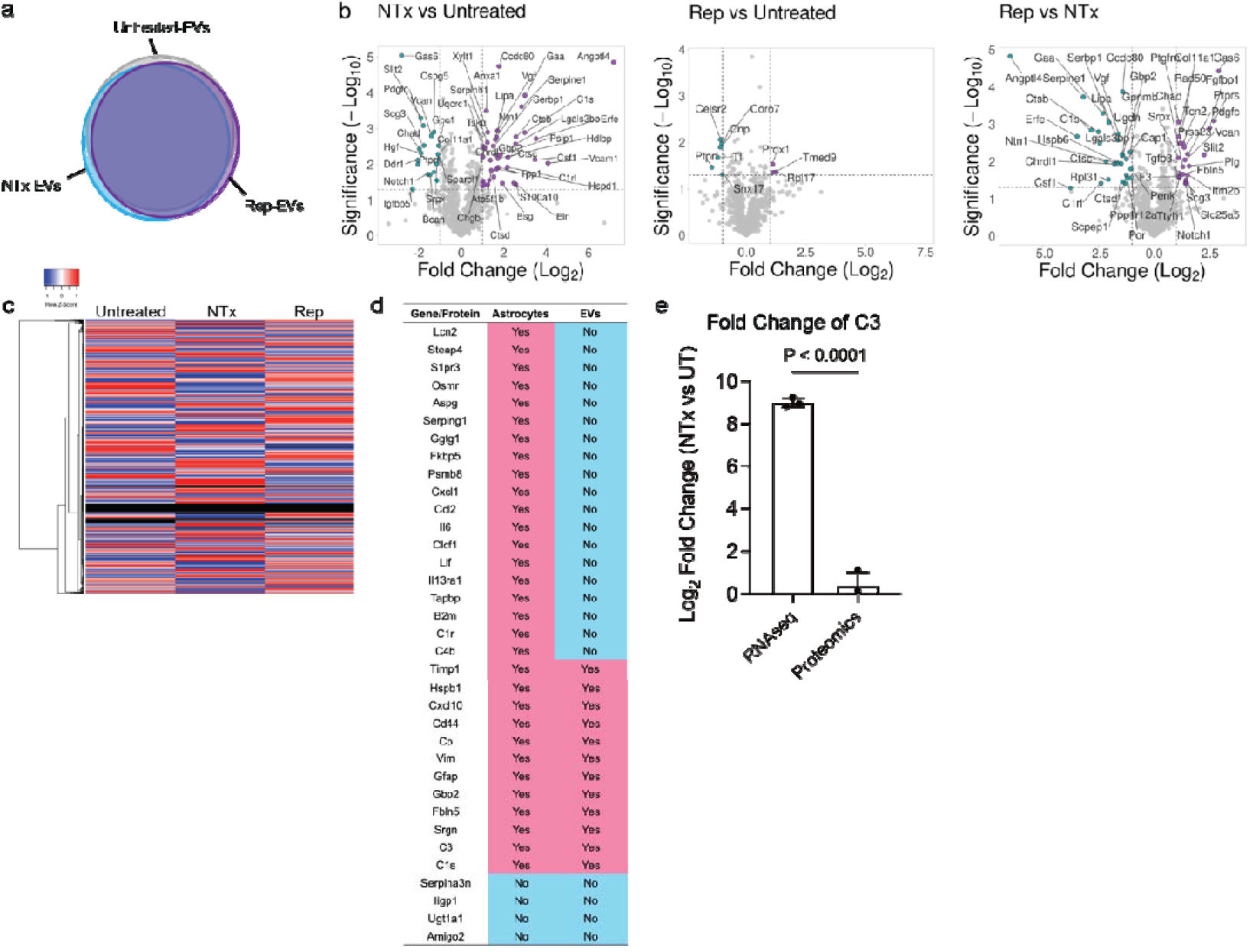
Inflammatory proteins are less presented in EVs especially for NTx astrocytes. a) Venn diagram showing the overlapping proteins among UT, NTx, and Rep astrocytes. b) Volcano plots with significance ratio of 1.3 and fold change of 1 showing differential expression of proteins between UT, NTx, and Rep-EVs. c) Heatmap of all proteins within UT, NTx, and Rep astrocytes. d) Comparison on the presence of reactive and/or inflammatory genes/proteins between astrocyte RNA sequencing and EV proteomics data. e) The difference of log2 fold change of C3 comparing NTx to UT between astrocyte RNA sequencing and EV proteomics data, showing inflammatory proteins were less encapsulated within EVs by astrocytes.

## References

1 Burda, J. E. & Sofroniew, M. V. Reactive gliosis and the multicellular response to CNS damage and disease. Neuron 81, 229–248 (2014).

2 Zamanian, J. L. et al. Genomic analysis of reactive astrogliosis. Journal of neuroscience 32, 6391–6410 (2012).

3 Williamson, M. R., Fuertes, C. J. A., Dunn, A. K., Drew, M. R. & Jones, T. A. Reactive astrocytes facilitate vascular repair and remodeling after stroke. Cell reports 35, 109048 (2021).

4 Llorente, I. L. et al. Patient-derived glial enriched progenitors repair functional deficits due to white matter stroke and vascular dementia in rodents. Science Translational Medicine 13 (2021).

5 Hu, S. et al. Exosome-eluting stents for vascular healing after ischaemic injury. Nature biomedical engineering 5, 1174–1188 (2021).

6 Hu, M. et al. Hepatic macrophages act as a central hub for relaxin-mediated alleviation of liver fibrosis. Nature nanotechnology 16, 466–477 (2021).

7 Li, L. et al. Transplantation of human mesenchymal stem-cell-derived exosomes immobilized in an adhesive hydrogel for effective treatment of spinal cord injury. Nano Letters 20, 4298–4305 (2020).

8 Safety of Mesenchymal Stem Cell Extracellular Vesicles (BM-MSC-EVs) for the Treatment of Burn Wounds, <https://www.clinicaltrials.gov/ct2/show/NCT05078385> (2021).

9 Chun, C. et al. Astrocyte-derived extracellular vesicles enhance the survival and electrophysiological function of human cortical neurons in vitro. Biomaterials 271, 120700 (2021).

10 You, Y. et al. Activated human astrocyte-derived extracellular vesicles modulate neuronal uptake, differentiation and firing. Journal of extracellular vesicles 9, 1706801 (2020).

11 Chaudhuri, A. D. et al. TNFα and IL-1ß modify the miRNA cargo of astrocyte shed extracellular vesicles to regulate neurotrophic signaling in neurons. Cell death & diseased, 1–8 (2018).

12 Liddelow, S. A. et al. Neurotoxic reactive astrocytes are induced by activated microglia. Nature 541, 481–487 (2017).

13 Sadtler, K. et al. Developing a pro-regenerative biomaterial scaffold microenvironment requires T helper 2 cells. Science 352, 366–370 (2016).

14 Guttenplan, K. A. et al. Neurotoxic reactive astrocytes induce cell death via saturated lipids. Nature 599, 102–107 (2021).

15 Wilson, K. L., Carmichael, S. T. & Segura, T. Injection of hydrogel biomaterial scaffolds to The brain after stroke. JoVE (Journal of Visualized Experiments), e61450 (2020).

16 Nih, L. R., Gojgini, S., Carmichael, S. T. & Segura, T. Dual-function injectable angiogenic biomaterial for the repair of brain tissue following stroke. Nature materials 17, 642–651 (2018).

17 Sideris, E. et al. Particle Hydrogels Decrease Cerebral Atrophy and Attenuate Astrocyte and Microglia/Macrophage Reactivity after Stroke. Advanced Therapeutics, 2200048 (2019).

18 Heras-Romero, Y. et al. Improved post-stroke spontaneous recovery by astrocytic extracellular vesicles. Molecular Therapy 30, 798–815 (2022).

19 Lenzini, S., Bargi, R., Chung, G. & Shin, J.-W. Matrix mechanics and water permeation regulate extracellular vesicle transport. Nature nanotechnology 15, 217–223 (2020).

20 Nih, L. R., Sideris, E., Carmichael, S. T. & Segura, T. Injection of microporous annealing particle (MAP) hydrogels in the stroke cavity reduces gliosis and inflammation and promotes NPC migration to the lesion. Advanced Materials 29, 1606471 (2017).

21 Griffin, D. R., Weaver, W. M., Scumpia, P. O., Di Carlo, D. & Segura, T. Accelerated wound healing by injectable microporous gel scaffolds assembled from annealed building blocks. Nature materials 14, 737–744 (2015).

22 Griffin, D. R. et al. Activating an adaptive immune response from a hydrogel scaffold imparts regenerative wound healing. Nature materials 20, 560–569 (2021).

23 Daly, A. C., Riley, L., Segura, T. & Burdick, J. A. Hydrogel microparticles for biomedical applications. Nature Reviews Materials 5, 20–43 (2020).

24 Lee, T. S., Kim, Y., Zhang, W., Song, I. H. & Tung, C.-H. Facile metabolic glycan labeling strategy for exosome tracking. Biochimica et Biophysica Acta (BBA)-General Subjects 1862, 1091–1100 (2018).

25 Baskin, J. M. et al. Copper-free click chemistry for dynamic in vivo imaging. Proceedings of the National Academy ofSciences 104, 16793–16797 (2007).

26 Nikolakopoulou, A. M. et al. Pericyte loss leads to circulatory failure and pleiotrophin depletion causing neuron loss. Nature Neuroscience 22, 1089–1098 (2019).

27 Kirby, L. et al. Oligodendrocyte precursor cells present antigen and are cytotoxic targets in inflammatory demyelination. Nature communications 10, 1–20 (2019).

28 Benowitz, L. I. & Carmichael, S. T. Promoting axonal rewiring to improve outcome after stroke. Neurobioiogy of diseases 28, 259–266 (2010).

29 Tong, L.-s. et al. Recombinant Gas6 augments Axl and facilitates immune restoration in an intracerebral hemorrhage mouse model. Journal of Cerebral Blood Flow & Metabolism 37, 1971–1981 (2017).

30 Takahashi, H. & Craig, A. M. Protein tyrosine phosphatases PTPδ, PTPσ, and LAR: presynaptic hubs for synapse organization. Trends in neurosciences 36, 522–534 (2013).

31 Taetzsch, T., Brayman, V. L. & Valdez, G. FGF binding proteins (FGFBPs): modulators of FGF signaling in the developing, adult, and stressed nervous system. Biochimica et Biophysica Acta (BBA)-Molecular Basis of Disease 1864, 2983–2991 (2018).

32 Sherchan, P. et al. Recombinant Slit2 reduces surgical brain injury induced blood brain barrier disruption via Robo4 dependent Rac1 activation in a rodent model. Scientific reports 7, 1–11 (2017).

33 Huang, Q. et al. Aligned graphene mesh-supported double network natural hydrogel conduit loaded with netrin-1 for peripheral nerve regeneration. ACS Applied Materials & Interfaces 13, 112–122 (2021).

34 Anderson, M. A. et al. Astrocyte scar formation aids central nervous system axon regeneration. Nature 532, 195–200 (2016).

35 Su, N. et al. Mesenchymal stromal exosome-functionalized scaffolds induce innate and adaptive immunomodulatory responses toward tissue repair. Science advances 7, eabf7207 (2021).

36 Jiang, Y. et al. Brain Microenvironment Responsive and Pro-angiogenic Extracellular Vesicle-Hydrogel for Promoting Neurobehavioral Recovery in Type 2 Diabetic Mice after Stroke. Advanced Healthcare Materials, 2201150 (2022).

37 Stogsdill, J. A. et al. Astrocytic neuroligins control astrocyte morphogenesis and synaptogenesis. Nature 551, 192–197 (2017).

